# New insights into the allosteric effects of CO_2_ and bicarbonate on crocodilian hemoglobin

**DOI:** 10.1101/2021.03.22.436447

**Authors:** Naim M. Bautista, Hans Malte, Chandrasekhar Natarajan, Tobias Wang, Jay F. Storz, Angela Fago

## Abstract

Crocodilians are unique among vertebrates in that their hemoglobin (Hb) O_2_ binding is allosterically regulated by bicarbonate, which forms in the red blood cell upon hydration of CO_2_. Although known for decades, this remarkable mode of allosteric control has not yet been experimentally verified with direct evidence of bicarbonate binding to crocodilian Hb, probably because of confounding CO_2_-mediated effects. Here we provide the first quantitative analysis of the separate allosteric effects of CO_2_ and bicarbonate on Hb of the spectacled caiman (*Caiman crocodilus*). Using thin-layer gas diffusion chamber and Tucker chamber techniques, we demonstrate that both CO_2_ and bicarbonate bind to Hb with high affinity and strongly decrease Hb-O_2_ saturation, and propose that both effectors bind to an unidentified positively charged site containing a reactive amino group in the low-O_2_ affinity T conformation of the Hb. These results provide the first experimental evidence that bicarbonate binds directly to crocodilian Hb and promotes O_2_ delivery independently of CO_2_. Using the gas-diffusion chamber, we observed similar effects in the Hbs of a phylogenetically diverse set of other caiman, alligator, and crocodile species, suggesting that the unique mode of allosteric regulation by CO_2_ and bicarbonate evolved >80-100 million years ago in the common ancestor of crocodilians. Taken together, our results show a tight and unusual linkage between O_2_ and CO_2_ transport in the blood of crocodilians, where build-up of blood CO_2_ and bicarbonate ions during breath-hold diving or digestion facilitates O_2_ delivery, while Hb desaturation facilitates CO_2_ transport as protein-bound CO_2_ and bicarbonate.

## Introduction

The oxygenation properties of vertebrate hemoglobins (Hbs) are sensitive to a number of small-molecule effectors present in the red blood cell (RBC), including protons, CO_2_, chloride ions, and organic phosphates. Allosteric regulation of Hb-O_2_ affinity by such effectors helps keep blood O_2_ supply at balance with tissue O_2_ consumption. These effectors bind preferentially to the low-affinity (deoxy) T-state of the Hb tetramer, thereby reducing Hb-O_2_ affinity by shifting the T-R equilibrium towards the T quaternary state (Perutz, 1976; Storz, 2019; Weber and Fago, 2004). Hence, increases in blood CO_2_ levels (as produced by metabolic activity) cause a reduction in Hb-O_2_ affinity, thereby facilitating O_2_ unloading to tissues. The underlying mechanism is the reaction of CO_2_ with N-terminal amino groups (-NH_2_) of the α- and β-chain subunits of the Hb tetramer (Kilmartin and Rossi-Bernardi, 1971), resulting in the formation of a carbamino product (-NH-COO^-^), which preferentially stabilizes the low-affinity T state of the Hb via interactions with positively-charged amino acid residues. CO_2_ also exerts an indirect allosteric effect on Hb-O_2_ affinity by reducing RBC pH. Upon entry in the RBC, CO_2_ reacts with water to form bicarbonate ions and protons, a reaction catalyzed by the enzyme carbonic anhydrase. Protons bind to Hb and promote O_2_ unloading via the Bohr effect, while bicarbonate ions leave the RBC in exchange for chloride ions.

Among vertebrate Hbs, those of crocodilians are reportedly unique in that their O_2_-affinity is allosterically regulated by binding of bicarbonate ions to the Hb protein (Bauer et al., 1981; Perutz et al., 1981). This unusual bicarbonate sensitivity of crocodilian Hb is thought to support aerobic metabolism when blood bicarbonate levels are elevated. This occurs during breath-hold diving (Bautista et al., 2021; Hicks and White, 1992), and during the postprandial ‘alkaline tide’ where gastric acid secretion causes a pronounced metabolic alkalosis (Busk et al., 2000).

The bicarbonate sensitivity of crocodilian Hbs was first described in the spectacled caiman (*Caiman crocodilus*) (Bauer et al., 1981; Perutz et al., 1981), but the allosteric effect was inferred without direct experimental evidence of bicarbonate binding to Hb. Subsequent studies on Hbs from other crocodilian species documented a direct allosteric effect of CO_2_ (Fago et al., 2020; Jensen et al., 1998; Weber et al., 2013), but the direct contribution of bicarbonate-binding has neither been confirmed in these studies nor quantified. The reason is likely that it is difficult to experimentally distinguish the allosteric effects of CO_2_ from those of bicarbonate and protons because at equilibrium all these three compounds are present in solution. Moreover, previous studies (Bauer et al., 1981) used different experimental and buffer conditions and rarely accounted for the potential effect of other anions, such as chloride ions, which are potent allosteric modulators of crocodilian Hb-O_2_ affinity (Fago et al., 2020; Storz, 2019; Weber et al., 2017).

Here, we use kinetic and equilibrium approaches at a constant pH and in the absence of chloride ions or organic phosphates to separate the allosteric effects of CO_2_ and bicarbonate on crocodilian Hb oxygenation. We provide a detailed functional analysis on the allosteric regulation of the Hb of the spectacled caiman, the same species that was originally studied by Bauer and coworkers (Bauer et al., 1981; Bauer and Jelkmann, 1977; Jelkmann and Bauer, 1980). We also confirm the same mode of allosteric regulation in the Hbs of 13 other caiman, alligator, and crocodile species, indicating that this unique property evolved in the common ancestor of extant crocodilians.

## Materials and Methods

### Animals and blood collection

Approximately 5 ml blood was drawn from the post-occipital sinus into heparinized syringes from three non-anesthetized adult spectacled caimans (*Caiman crocodilus*; Linnaeus 1758; around 1.5 kg each) of undetermined sex. The procedure lasted less than 5 min. Blood was immediately centrifuged at 4°C (5000 g, 5 min) to remove plasma and RBCs were then washed five times in 0.9 % NaCl, 0.5 mM EDTA and stored at −80 °C until further processing. As described previously (Fago et al., 2020), we sampled blood and prepared hemolysates from 13 other crocodilian species (American alligator, *Alligator mississippiensis,* [Duadin, 1802]; Chinese alligator, *A. sinensis,* [Fauvel, 1879]; broad-snouted caiman, *Caiman latirostris* [Duadin, 1802]; yacaré caiman, *C. yacare* [Duadin, 1802]; black caiman, *Melanosuchus niger,* [Spix, 1825]; smooth-fronted caiman, *Paleosuchus trigonatus,* [Schneider, 1801]; American crocodile, *Crocodylus acutus,* [Cuvier, 1807]; Philippine crocodile, *C. mindorensis* [Schmidt, 1928]; Nile crocodile, *C. niloticus* [Laurenti, 1768]; New Guinea crocodile, *C. novaeguineae* [Schdmit, 1928]; saltwater crocodile, *C. porosus* [Schneider, 1801]; Cuban crocodile, *C. rhombifer* [Cuvier, 1807]; Siamese crocodile, *C. siamensis* [Schneider, 1801]).

All protocols of animal handling and sampling were performed in accordance with the Danish Law for Animal Experimentation and were approved by the Danish Animal Experiments Inspectorate, under permit number 2018-15-0201-01507.

### Preparation of the hemolysate

For each individual spectacled caiman (N=3), the hemolysate was prepared by dilution (1:5) of washed RBCs in chloride-free 10 mM HEPES, pH 7.6, 0.5 mM EDTA, and incubated on ice for 30 min. The hemolysate was then centrifuged at 4°C (12000 g, 20 min) to remove cell debris and then passed (‘stripped’) on a PD-10 desalting column (GE Healthcare) equilibrated with 10 mM HEPES, pH 7.6, 0.5 mM EDTA to remove small molecular weight molecules, including Hb’s allosteric effectors. Chloride-free HEPES buffers were used throughout. Absence of chloride in solutions was verified by model 926S Mark II chloride analyzer (Sherwood Scientific Ltd, Cambridge, UK).

### Discriminating the effects of CO_2_ and bicarbonate

To discriminate between the effect of CO_2_ and bicarbonate on Hb oxygenation, we measured the time-course of changes in O_2_ saturation of stripped hemolysate and purified Hb (see Hemoglobin purification) after addition and removal of 1% CO_2_ gas at a fixed PO_2_. These experiments were performed in the presence and absence of 1 mM acetazolamide, a specific inhibitor of carbonic anhydrase contained in the hemolysate, in order to manipulate the rate of bicarbonate formation from CO_2_ (Fago et al., 1999). Experiments were performed at 25°C using a thin-layer modified diffusion chamber technique (Cadiz et al., 2019; Natarajan et al., 2018; Storz et al., 2020; Weber et al., 2017). In brief, 4 µL samples were placed in the temperature-controlled gas diffusion chamber connected to a Cary 60 UV-Vis spectrophotometer equipped with fibre optic probes (Agilent Technologies, CA, USA), and O_2_ saturation was determined from the absorption trace at 415 nm. Discrete values of PO_2_ and PCO_2_ within the chamber were obtained by mixing O_2_ and CO_2_ with ultrapure N_2_ gas using a programmable Gas Mixing System (GMS – Loligo Systems, Viborg, Denmark), with the gas mixture constantly flowing above the sample surface. The experiments were performed at a fixed PO_2_ corresponding approximately to 50% O_2_ saturation, under the same conditions as our previous study on crocodilian Hbs (0.3 mM heme, 0.1 M HEPES, pH 7.2, 0.5 mm EDTA) (Fago et al., 2020). Heme concentration of samples was measured spectrophotometrically using published extinction coefficients at 577 and 542 nm (Van Assendelft, 1970).

Throughout this study, the pH of Hb samples equilibrated with the same CO_2_ of the experiments was routinely measured using a pH electrode InLab® Micro (Mettler Toledo) and it did not change appreciably (range 7.16-7.21).

### Measuring O_2_ equilibrium curves

We measured O_2_ equilibrium curves of stripped hemolysate and purified Hb (see Hemoglobin purification) using the same diffusion chamber technique described above at 25 °C under identical buffer conditions (0.3 mM heme, 0.1 M HEPES, pH 7.2, 0.5 mm EDTA). O_2_ affinity (P_50_, the PO_2_ at 50% O_2_ saturation) and Hill’s cooperativity coefficient (n) were calculated by non-linear regression fitting the Hill equation (Eq. 1) to the saturation data (Janecka et al., 2015; Jendroszek et al., 2018)

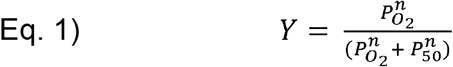

where Y is the O_2_ saturation and PO_2_ is the O_2_ tension.

### Hemoglobin purification and mass spectrometry analysis

Hb was purified from endogenous carbonic anhydrase by gel filtration using fast protein liquid chromatography (FPLC) on an ÄKTA Pure Protein Purification System equipped with a Superdex 75 10/300 GL gel-filtration column and a 100 µL loop (GE Healthcare Life Sciences). Buffer was 10 mM HEPES, pH 7.6, 0.5 mM EDTA, 0.15 M NaCl, and flow rate was 0.5 ml/min. Absorbance was measured at 280 nm. The column was calibrated with standard proteins (Sigma Aldrich) of known molecular mass: horse myoglobin (17.6 kDa), bovine carbonic anhydrase II (30 kDa) and human adult Hb (64.6 kDa). The Hb fractions purified by gel filtration were concentrated by ultracentrifugation and dialyzed against 10 mM HEPES, pH 7.5, 0.5 mM EDTA to remove NaCl. After dialysis, absence of Cl^-^ in the samples was confirmed by using a MK II Chloride Analyzer 926S (Sherwood Scientific Ltd, Cambridge, Uk). Hb fractions were analysed for the presence or absence of intermolecular disulfide bonds by SDS polyacrylamide gel electrophoresis (range 4-12 %) using NuPAGE® Novex® Bis-Tris Pre-Cast Gels and stained with Coomassie blue, in the presence and absence of the reducing agent DTT.

To establish whether carbonic anhydrase (~30 kDa) co-eluted with Hb on FPLC, we ran the purified Hb fractions on a mini-protean precast 4-20% SDS PAGE gel (Bio-Rad, Hercules, CA), which we then stained with Coomassie brilliant blue-G. We conducted tandem mass spectrometry (MS/MS) analyses on the bands ranging from ~25 to 50 kDa in size (Fig.S3). The stained bands were excised and processed for in-gel tryptic digestion (Shevchenko et al., 2006) and the eluted peptides were then analyzed using a Thermo Orbitrap Fusion Lumos Tribrid (Thermo Scientific™) mass spectrometer in data-dependent acquisition mode. Peptides were identified by searching MS/MS data against the UniprotKB/Swiss-Prot database. The search was set up for full tryptic peptides with a maximum of two missed cleavage sites. The precursor mass tolerance threshold was set as 10 ppm and maximum fragment mass error was set at 0.02 Da. Qualitative analysis was performed using PEAKS X software. The significance threshold of the ion score was calculated based on a false discovery rate of ≤ 1%.

### CO_2_ effect on O_2_ affinity and CO_2_ saturation curves

To establish whether the purified Hb fractions had different sensitivity to CO_2_, we determined O_2_ equilibrium curves for each Hb fraction at 25 °C (0.3 mM heme, 0.1 M HEPES, pH 7.2, 0.5 mm EDTA) at discrete CO_2_ tensions in the gas mixture, and calculated P_50_ and n at each PCO_2_ as described above (see Measuring O_2_ equilibrium curves).

For each purified Hb fraction, we determined apparent CO_2_ saturation curves by measuring the decrease in the O_2_ saturation upon addition of CO_2_ in the gas mixture, while keeping a fixed PO_2_ (approximately corresponding to the P_50_ of that Hb fraction in the absence of CO_2_). A new Hb sample (4 µL) was used for measurements at each given PCO_2_. Experiments were performed using the modified diffusion chamber technique at 25 °C, 0.3 mM heme, 0.1 M HEPES, pH 7.2, 0.5 mm EDTA. Apparent CO_2_ saturation curves for each purified Hb were obtained by non-linear regression fitting of a single hyperbola (Eq. 2) to the data

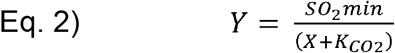

where Y is O_2_ saturation, SO_2min_ is the O_2_ saturation at saturating PCO_2_ values (plateau) and X is PCO_2._ This analysis provided the apparent affinity constant for the allosteric binding of CO_2_ to the Hb (K_CO2_, the PCO_2_ at half-CO_2_ saturation). We also examined the possibility of two binding sites by fitting a double hyperbola equation to the data (see Supplementary material Table S1).

### Bicarbonate effect on O_2_ release and bicarbonate saturation curves

To determine the effect of bicarbonate on Hb oxygenation, we equilibrated samples of each purified Hb fraction (0.05 mM heme, 0.1 M HEPES buffer pH 7.2, 0.5 EDTA) in Eschweiler tonometers for at least 40 min at 25°C with a gas mixture of 2% air and 98% N_2_ generated by a gas mixing pump (Wösthoff, Bochum, Germany). The same gas mixture (corresponding to a PO_2_ close to the P_50_ of the Hb) was also used to equilibrate a freshly prepared stock solution of sodium bicarbonate (NaHCO_3_) dissolved in MilliQ H_2_O in a separate tonometer at the same temperature. After equilibration, the Hb sample was transferred with a gastight syringe into a 2.5 ml Tucker chamber (Tucker, 1967), previously flushed with 2% air and 98% N_2._ The chamber was equipped with a fibre optic O_2_ probe connected to a FireStingO2 (FSO2-1) oxygen and temperature meter and controlled by FireSting logger V.2.365 software for PO_2_ signal recording (PyroScience, Germany). Following stabilization of the PO_2_ signal of the Hb sample in the chamber, a small volume (< 60 µL) of the NaHCO_3_ solution was injected into the Tucker chamber to achieve discrete NaHCO_3_ concentrations (2, 4, 7.7, 15.4, 23.1 and 30 mM), and the subsequent increase in PO_2_ in the chamber (due to bicarbonate-induced O_2_ unloading by the Hb) was recorded. The bicarbonate concentrations of the stock solutions were 0.5, 1, 1.2, or 1.3 M, chosen based on the desired final concentration within the chamber, to maintain a similar injection volume (< 60 µl) in all samples. A new Hb sample was used for each bicarbonate addition and separate experiments were performed for the Hb fractions of each of the three individuals. Two-point calibration of the oxygen probe was performed immediately before recording each trace by flushing the chamber with water-saturated air or N_2_ and taking into accounts the barometric pressure on the day. The pH of the bicarbonate-containing Hb solution was measured at the end of each experiment using a Mettler Toledo pH electrode InLab® Micro and no significant change was observed (range 7.19-7.23).

Apparent bicarbonate saturation curves for each purified Hb were obtained by non-linear regression fitting of a single hyperbola (Eq. 3) to the data

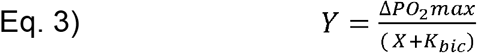

where Y is ∆PO_2_, ∆PO_2_max is the ∆PO_2_ at saturating bicarbonate concentrations (plateau) and X is [NaHCO_3_] in mM. This analysis provided the apparent affinity constant for the allosteric binding of bicarbonate to the Hb (K_bic_, the bicarbonate concentration at half-bicarbonate saturation).

### Statistical analysis

Statistical analysis and curve fitting were performed using SigmaPlot version 12 (SigmaStat) and Prism V.8 (GraphPad). The obtained parameters SO_2min_ and K_CO2_ were statistically compared by two-way ANOVA considering CO_2_ and Hb fraction as factors. The K_CO2_ values obtained for the Hb fractions were statistically compared by a t-test analysis. Similarly, the ∆PO_2_max and K_bic_ values obtained for the Hb fractions were statistically compared by t-test. Statistical significance was considered at α < 0.05. Data is presented as mean ± s.e.m unless otherwise stated.

## Results and Discussion

### Reversible effects of CO_2_ on hemolysate oxygenation

To investigate the mode of allosteric regulation of crocodilian Hb, we first tested whether the CO_2_ effect on Hb-O_2_ saturation could be attributed exclusively to bicarbonate, as originally proposed by Bauer and coworkers (Bauer et al., 1981). In these experiments, we used hemolysates of spectacled caiman (N=3 individuals) and 13 other crocodilian species representing the two most speciose families, Alligatoridae and Crocodylidae, which together capture the deepest evolutionary split among all living crocodilians. We used a kinetic approach that we originally developed to unravel the allosteric effect of bicarbonate in hagfish Hb, which is insensitive to CO_2_ (Fago et al., 1999). This method takes advantage of active endogenous carbonic anhydrase in the hemolysate to rapidly convert added CO_2_ to bicarbonate, as well as the carbonic anhydrase inhibitor acetazolamide to specifically inhibit this fast enzymatic reaction. Thus, upon addition of acetazolamide, a slower effect of CO_2_ on O_2_ saturation indicates that CO_2_ must convert to bicarbonate (via the slow non-catalysed reaction) to be able to affect oxygenation and that bicarbonate – not CO_2_ – is the true allosteric effector (Fago et al.,1999). As shown in Figure 1, at a fixed PO_2_ (yielding ~50% O_2_ saturation), addition of 1% CO_2_ in the gas mixture caused a rapid and strong decrease in the O_2_ saturation down to ~10 % in the stripped hemolysate of spectacled caiman (Fig. 1A). We also observed the same response in the hemolysates of other crocodilian species (Fig. S1). This CO_2_-mediated decrease in O_2_ saturation was fully reversible, and 50% saturation was restored when CO_2_ was removed from the gas mixture (Fig. 1A). The decrease in O_2_ saturation was equally fast when carbonic anhydrase present in the hemolysate was inhibited by acetazolamide (Fig. 1B, Fig S1). This result indicates that CO_2_ binds to the Hb directly without being converted to bicarbonate and that CO_2_ is a potent allosteric effector of crocodilian Hbs in its own right. This is not consistent with results reported by Bauer and coworkers, which were obtained under different experimental conditions, perhaps due to the use of chloride-containing solutions (Bauer et al., 1981). Also, the fact that the decrease in O_2_ saturation was of the same magnitude in the presence and absence of acetazolamide (Fig. 1, Fig.S1) indicates that there is no additional binding of bicarbonate (which would form rapidly without acetazolamide) elsewhere on the protein. Thus, the proposed allosteric effect of bicarbonate must occur via binding of bicarbonate to the same protein site as CO_2_.

**Figure 1.**
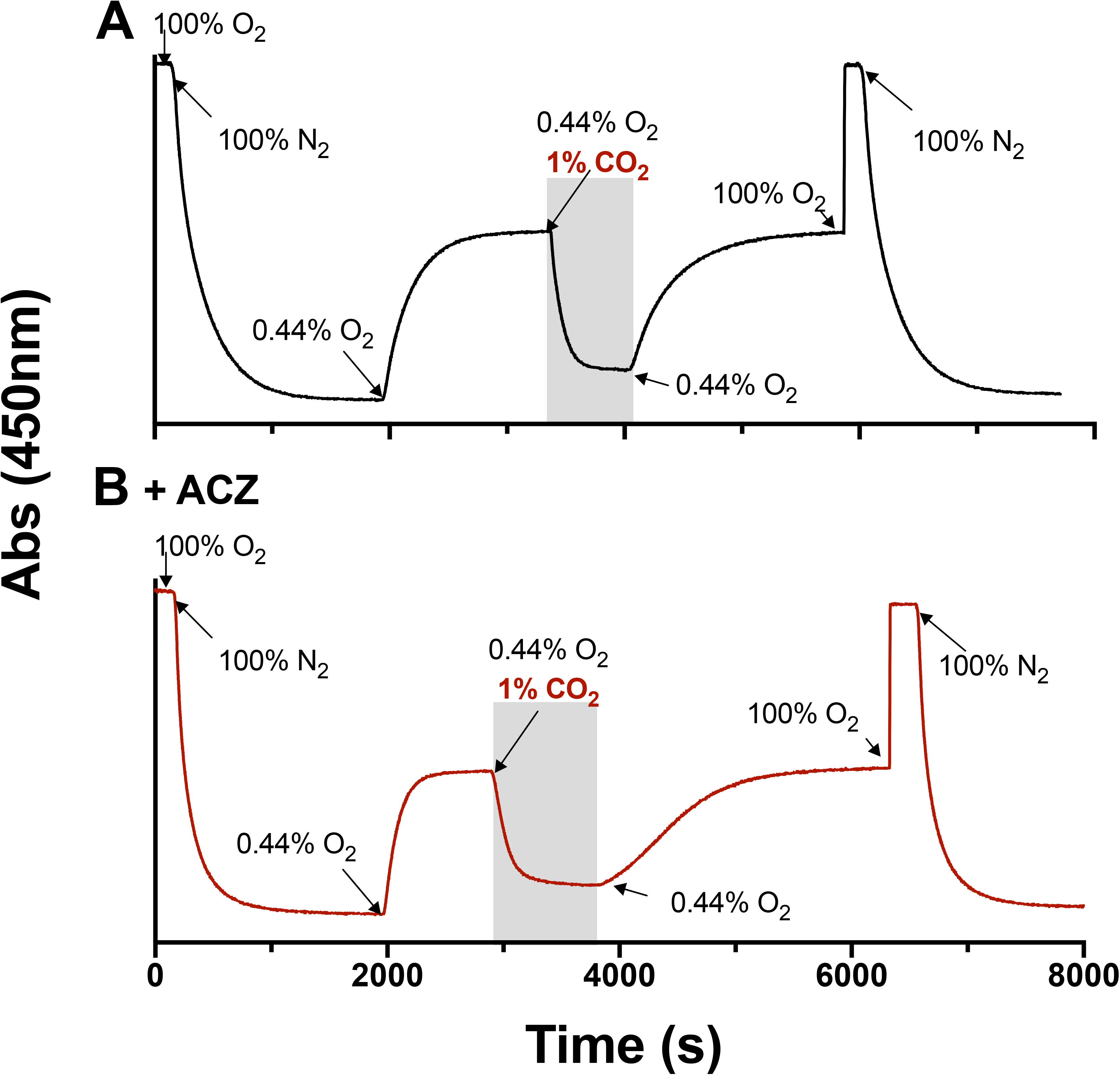
Reversibility of CO_2_ effect on O_2_ saturation in the hemolysate of the spectacled caiman. Representative traces in the absence **A)**and presence **B)**of acetazolamide (ACZ) monitored by the diffusion chamber technique. After equilibration with 100% O_2_ (100% O_2_ saturation) and 100% N_2_ (0% O_2_ saturation), the hemolysate was equilibrated with 0.44% O_2_ to achieve approximately 50% O_2_ saturation before equilibration with 1% CO_2_, 0.44% O_2_, as highlighted by the shaded area. Changes in the % gas in the mixture are indicated by arrows. Traces are representative of N=3 biological replicates.

The fact that the recovery to ~50% O_2_ saturation upon removal of CO_2_ was slower when carbonic anhydrase was inhibited (Fig. 1, Fig. S1) could indicate that carbonic anhydrase was somehow involved in the dissociation of Hb-bound CO_2_. The addition of bovine carbonic anhydrase II to spectacled caiman hemolysate produced similar traces as those of Figure 1 (Fig. S2), indicating that endogenous carbonic anhydrase was indeed present in the hemolysate.

### Functionally identical Hb fractions differing in molecular mass

In order to determine whether endogenous carbonic anhydrase was involved in the reversibility of CO_2_-mediated responses on Hb oxygenation, we passed spectacled caiman hemolysate through a gel-filtration column to remove carbonic anhydrase. Spectacled caiman expresses a single adult Hb isoform (Jelkmann and Bauer, 1980), similar to all other crocodilians examined (Fago et al., 2020; Hoffmann et al., 2018). Results from gel filtration showed differences in the apparent molecular mass of the Hb, with part of the Hb assembled into a higher-order aggregate – presumably an octamer – with an apparent mass of 95.7 kDa, while most Hb was present as a regular tetramer (Fig. 2A, B), as also reported earlier in the same species (Bauer and Jelkmann, 1977). We named these Hb fractions Hb1 and Hb2, respectively. Analysis of Hb1 and Hb2 by SDS-PAGE under reducing (Fig. 2C) and non-reducing conditions (Fig. S3) showed identical patterns and intensity of bands indicating that high-order aggregates are not formed via disulfide bonds as in the Hb aggregates of other crocodilians (Weber et al., 2013) and non-avian reptiles (Petersen et al., 2017; Storz et al., 2015).

**Figure 2.**
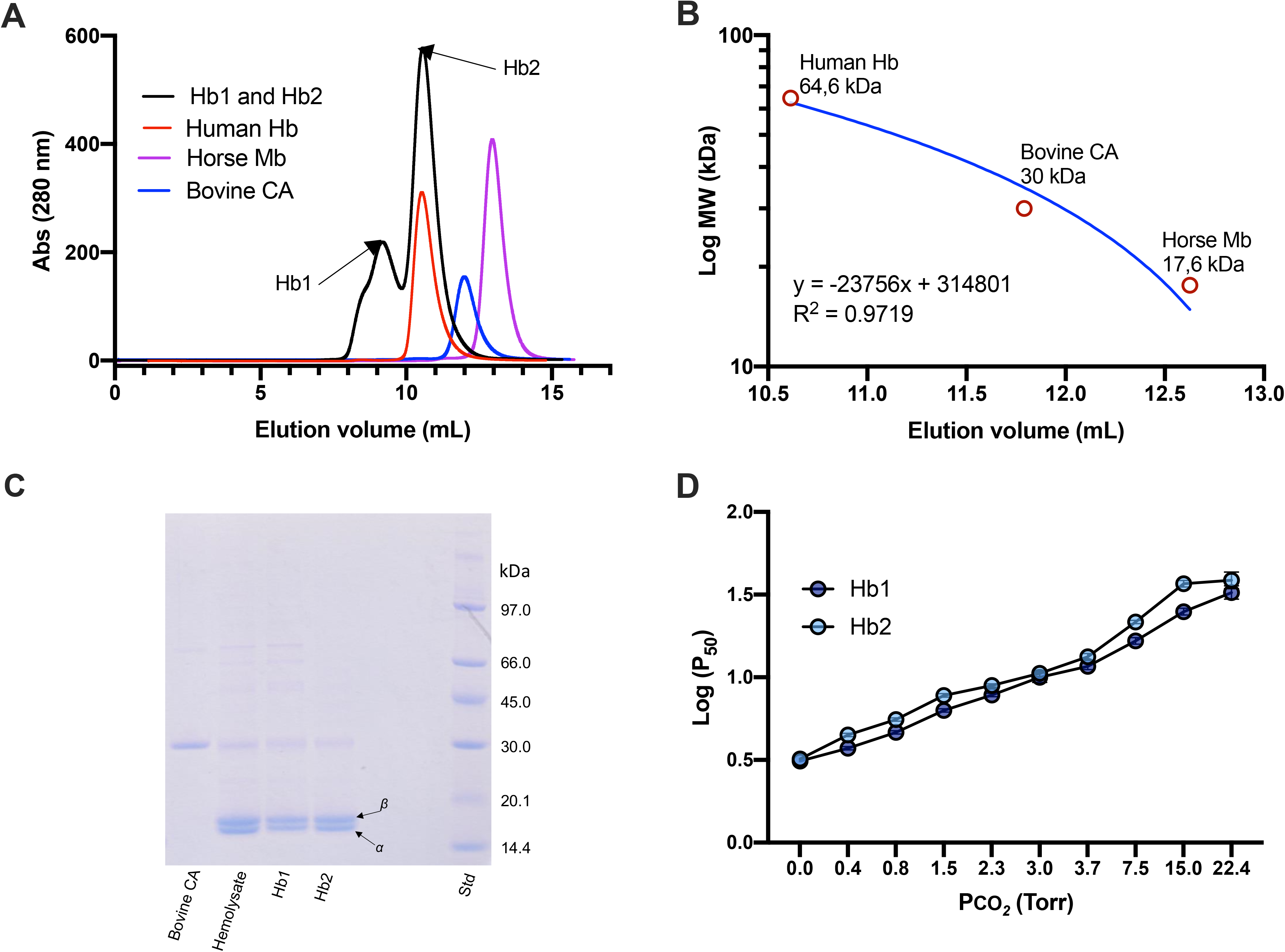
FPLC purification, SDS-PAGE and functional analysis of Hb fractions of spectacled caiman. **A)**Elution profile of caiman hemolysate (representative of N=3 biological replicates), resolving in two peaks (Hb1 and Hb2) indicated by arrows, is shown superimposed to that of standard proteins with known molecular mass human Hb (64.6 KDa), horse myoglobin (Mb, 17.6 KDa) and bovine carbonic anhydrase II (CA, 30 KDa). **B)**Calibration curve by regression analysis. **C)**SDS PAGE under reducing conditions (DTT 10 mM) of (lanes from left to right) bovine CA II, spectacled caiman hemolysate, purified Hb1, purified Hb2 and standard protein (STD) as indicated. The bands corresponding to α and β-type chains in the hemolysate and in the two purified Hb fractions are indicated. **D)**logP_50_ measured at various PCO_2_ values, showing identical effect of CO_2_ on the O_2_ affinity of purified Hb1 and Hb2 fractions (N=3).

SDS-PAGE of Hb1 and Hb2 revealed not only the α- and β-type Hb chains, but also less intense bands of higher molecular mass that could correspond to carbonic anhydrase, having a molecular mass of 30 kDa (Fig. 2C). The identity of these bands, corresponding to a size range of ~25 to 50 kDa, was established by mass spectrometry. The ~30 kDa band corresponds to stable intact heterodimers (Hb α and β-chains), possibly associated with aggregate formation of the Hb, and the ~42 kDa band corresponds to actin protein. Thus, carbonic anhydrase was effectively removed from both Hb1 and Hb2.

As a next step, we investigated the reversibility of CO_2_ binding on O_2_ saturation for the purified Hb1 and Hb2 fractions in the absence and presence of acetazolamide (Fig. S4). As for the hemolysate (Fig. 1), return to the original O_2_ saturation upon removal of CO_2_ was slower in the presence of acetazolamide. This result suggests that acetazolamide somehow interferes with Hb-CO_2_ dissociation, but not with Hb-CO_2_ binding.

To establish whether purified Hb1 and Hb2 had different O_2_ binding affinities and CO_2_ sensitivities, we measured O_2_ equilibrium curves under identical buffer conditions at different PCO_2_ values. These experiments yielded identical P_50_ at a range of PCO_2_ values for both Hb fractions (Fig. 2D) and highly similar cooperativity coefficients (Fig. S5), indicating that assembly of the Hb into higher-order aggregates did not alter oxygenation properties and CO_2_ sensitivities.

### CO_2_ apparent binding curves and affinity of Hb fractions

We used the diffusion chamber technique to determine the apparent affinity for CO_2_ of the purified Hb1 and Hb2 fractions (Fig. 3A-C). These experiments measured the decrease in O_2_ saturation at different PCO_2_ values, while maintaining a fixed PO_2_ roughly corresponding to the P_50_ of each Hb measured in the absence of CO_2_ (Fig. 3A). The Hb1 and Hb2 fractions exhibited identical changes in O_2_ saturations upon additions of CO_2_ (two-way ANOVA, P = 0.543), and data were therefore pooled. These data showed a single hyperbolic relationship (Fig. 3B,C), indicating the presence of a single type of binding site for CO_2_ in each Hb with apparent affinities (expressed as PCO_2_) corresponding to 1.48 ± 0.22 and 1.32 ± 0.15 torr for Hb1 and Hb2, respectively, that were not significantly different from each other (t-test, P = 0.570). The use of a double hyperbola equation did not narrow the confidence interval of the fitted parameters (Table S1), so the data support the existence of a single binding site for CO_2_. For each Hb fraction, the apparent affinity for CO_2_ is well below PCO_2_ values reported for crocodilian blood of ~29-36 torr during digestion (Busk et al., 2000), and 16.8 torr at rest and 26.7 torr during diving (Bautista et al., 2021), suggesting that crocodilian Hb would be largely saturated with CO_2_ under physiological conditions.

**Figure 3.**
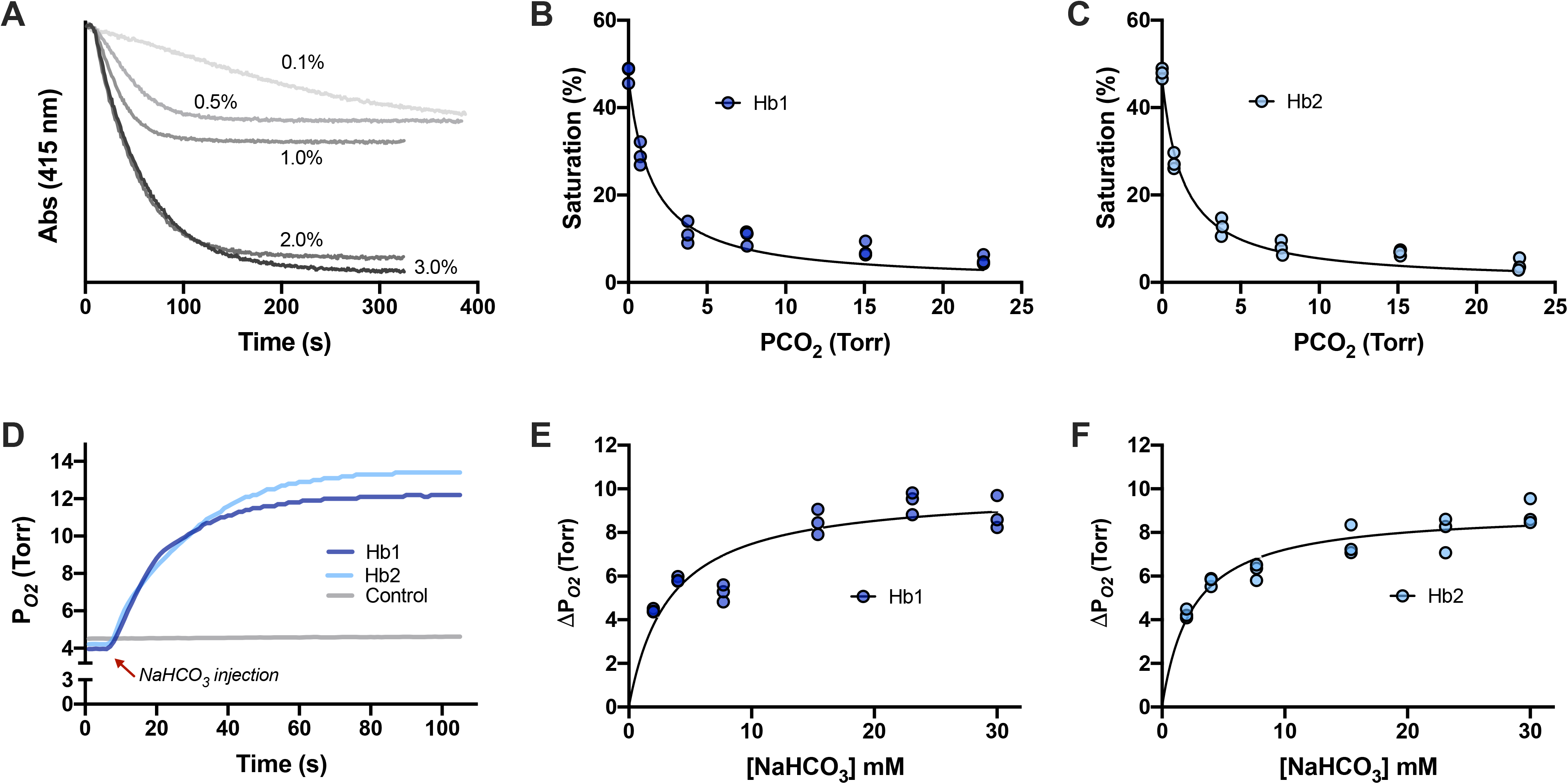
Effect of CO_2_ and bicarbonate on the O_2_ saturation of purified Hb1 and Hb2 from spectacled caiman and apparent CO_2_ and bicarbonate binding curves. **A)**Representative traces measured using the thin-layer diffusion chamber of the different decreases in Hb saturation induced by changes in PCO_2_ at a fixed PO_2_ close to P_50_ of 0.44 % O_2_ (PO_2_ ~3.3 Torr). Apparent CO_2_ saturation curves of **B)**Hb1 and **C)**Hb2 purified from 3 individuals obtained as shown in panel **A)**. Non-linear regression fitting of a single hyperbola equation is shown (details and fitted parameters are in Supplementary material Table S1). **D)**Representative traces showing the increase in O_2_ released from Hb1 (dark blue) and Hb2 (light blue) upon injection of bicarbonate at a fixed PO_2_ of ~4.2 torr. The gray line represents a control trace in which bicarbonate was added to buffer alone. Apparent bicarbonate saturation curves of **E)**Hb1 and **F)**Hb2 obtained as shown in panel **D)**. Non-linear regression fitting of a single hyperbola is shown (details and fitted parameters are in Supplementary material Table S2).

### Bicarbonate apparent binding curves and affinities of Hb fractions

The results of Figure 1 suggest that bicarbonate may allosterically bind to the same protein site as CO_2_. To confirm experimentally the direct allosteric effect of bicarbonate, we used purified Hb1 and Hb2 fractions of spectacled caiman in a Tucker chamber in the absence of CO_2_. In these experiments, we kept the Hb solution at a fixed PO_2_ (approximately corresponding to the P_50_) and measured the increase in PO_2_ following the injection of a measured small volume of bicarbonate stock solution (Fig. 3D-F). The results of these experiments show that bicarbonate had a dose-dependent effect on the O_2_ released from the Hb following a simple hyperbolic function, indicating the existence of a single type of allosteric binding site. Injections of bicarbonate solutions in buffer did not increase PO_2_ in the chamber (Fig. 3D). From the bicarbonate binding curves, the apparent bicarbonate affinities were 3.31 ± 0.35 and 2.48 ± 0.16 mM for Hb1 and Hb2, respectively (Table S2), which are not significantly different from each other (P = 0.094). These values are below the range of bicarbonate concentrations measured in crocodilian arterial blood of ~24-36 mM (from resting to digesting) (Busk et al., 2000; Jensen et al., 1998) and 11.4 mM at rest and 16.8 mM during diving in erythrocytes (Bautista et al., 2021), suggesting that crocodilian Hb is fully saturated with bicarbonate under physiological conditions. These results provide the first experimental evidence that bicarbonate binds directly to crocodilian Hb and promotes O_2_ delivery, independently of CO_2_.

Interestingly, the affinity for bicarbonate of spectacled caiman Hb in the deoxy state reported by Bauer and coworkers (Bauer et al., 1981) was ~0.7 mM, close to our estimated values at the same pH and temperature.

### A structural model for CO_2_ and bicarbonate allosteric binding

In analogy with other Hbs, we propose that CO_2_ chemically binds to the crocodilian Hb protein to form a carbamino compound (-NH-COO^-^) at a non-protonated amino group (-NH_2_) (see reaction scheme in Fig. 4A). The negatively charged carbamino group would then be stabilized by nearby positively charged amino acid residues on a specific T-state binding site (Fig. 4B), thereby causing a decrease in O_2_ affinity. Since pH does not change upon addition of CO_2_, as reported here and earlier (Jensen et al., 1998), the proton generated in carbamino formation (Fig. 4A) is picked up by the protein molecule, likely contributing to further electrostatic stabilization of the negative carbamino derivative. We note that there is very little structural difference between covalent CO_2_ binding via carbamino formation and non-covalent bicarbonate binding to the Hb protein (Fig. 4B), consistent with the inference from our results (Fig. 1) that both allosteric effectors bind to the same site.

**Figure 4.**
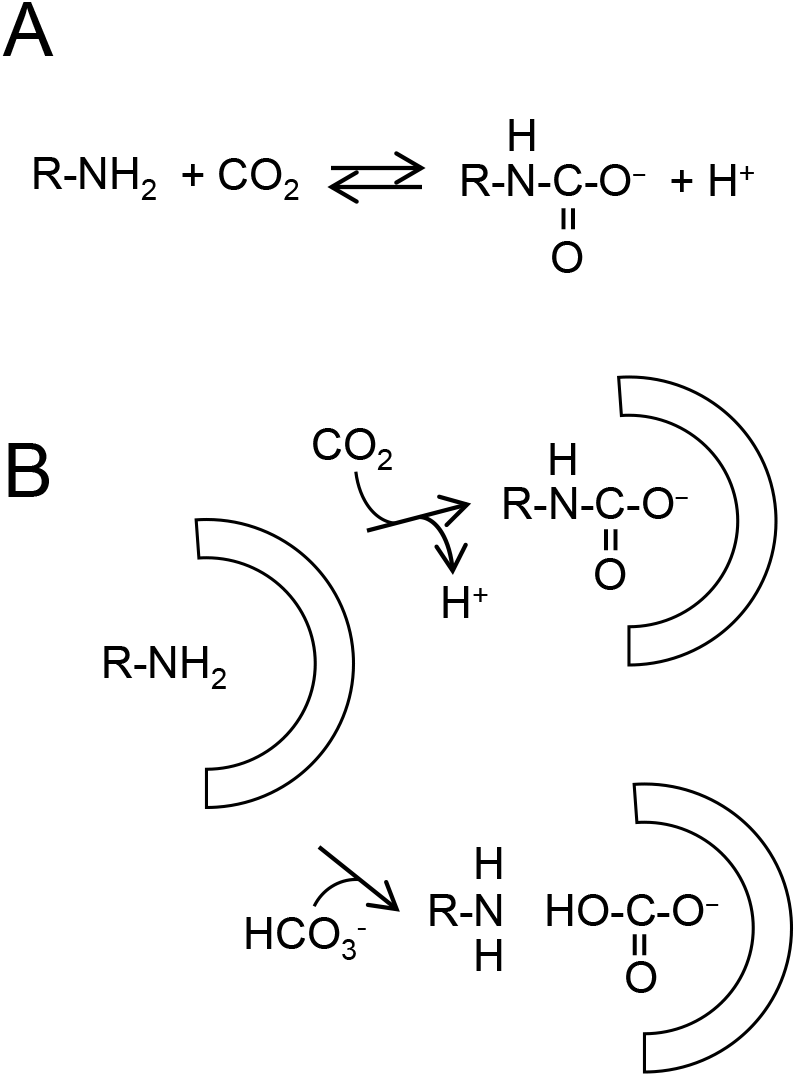
Proposed mechanistic model of the observed CO_2_ and bicarbonate dependent changes in Hb oxygenation in crocodilians. **A)**General scheme of carbamino protein derivative formation (R-NH-COO^-^) at a non-protonated amino group (R-NH_2_) from CO_2_. **B)**Identical putative binding site for CO_2_ (top) and bicarbonate (bottom) on the T-state of crocodilian Hb, including a reactive uncharged amino group surrounded by positively charged residues (indicated by the half circle). The negatively charged carbamino formed at the amino group (top) is stabilized by the positively charged amino acid residues, stabilizing the T state and causing a decrease in O_2_ saturation. The proton generated in carbamino formation is picked up by the protein molecule, likely contributing to further electrostatic stabilization of the carbamino. Note the very little structural difference between covalent CO_2_ binding via carbamino formation (top) and non-covalent bicarbonate binding (bottom) to the protein.

As also concluded in previous studies (Bauer et al., 1981; Perutz et al., 1981), our results (Fig. 3) and the two-fold symmetry of the tetrameric Hb molecule indicate the existence of two identical binding sites for CO_2_ and bicarbonate per tetramer in the T state of crocodilian Hb, each including a reactive non-protonated amino group close to positively charged residues. Carbamino formation in human Hb occurs at the α and β subunit N-termini (Bauer and Schröder, 1972; Kilmartin and Rossi-Bernardi, 1971; Perrella et al., 1975), but in crocodilian Hbs, these groups are acetylated to a variable extent without affecting the strong sensitivity to CO_2_ (Fago et al., 2020; Natarajan et al., 2020). In the original structural model proposed by Perutz and coworkers (Perutz et al., 1981) two bicarbonate ions bind two identical sites lining the central cavity of T-state crocodilian Hb, where organic phosphates normally bind: Lys82β, Glu144β, and the N-terminus of the α subunit. However, the smooth-fronted caiman (*Paleosuchus trigonatus*) possesses Gln at 82β rather than positively charged Lys (Fago et al 2020), and yet the Hb of this species exhibits the same CO_2_/bicarbonate effect as that of other crocodilians (Fig. S1). Moreover, acetylation of the α-chain N-termini of crocodilian Hbs does not appear to affect CO_2_ sensitivity (Natarajan et al. 2020). Based on site-directed mutagenesis of recombinant human Hb, other studies (Komiyama et al., 1996; Komiyama et al., 1995) have proposed a more complex mechanism, involving numerous residues at the α_1_β_2_ interface of the Hb tetramer, in particular, Lys38β, which is present in all crocodilian Hbs. Molecular modeling has revealed numerous positively charged residues in the central cavity of alligator Hb, including Arg135β involved in several interactions (Fago et al., 2020). In the vicinity of other positively charged residues, the amino group of Lys or Arg side chains, which normally have high pKa values and are protonated, would be largely in the non-protonated form at physiological pH values and would therefore be available for carbamino formation. Mutagenesis studies using recombinant crocodile Hb, rather human Hb, are underway in our lab to test these possibilities. Although the residues involved remain to be identified, it appears from our study that there is great diversity among chordate Hbs in the allosteric control of Hb function by CO_2_ and bicarbonate, with Hbs sensitive to only CO_2_ (e.g. human Hb), only bicarbonate (hagfish), or both in combination (crocodile).

## Conclusions

Our results indicate that the O_2_-affinity of crocodilian Hb is strongly modulated by allosteric binding of both CO_2_ and bicarbonate. Both effectors bind with high affinity at the same positively charged site located in the T-state and involving a reactive amino group, possibly the side chain of Lys38β, on each half-tetramer. Thus, it appears that the evolution of a particularly strong allosteric effect of CO_2_ in crocodilian Hbs would make these Hbs also sensitive to bicarbonate ions, whereas bicarbonate sensitivity apparently does not necessarily imply CO_2_ sensitivity, as it is the case for hagfish Hbs. Physiologically, the strong Hb sensitivities to CO_2_ and bicarbonate provides the basis for a unique linkage between O_2_ and CO_2_ transport in the blood of crocodilians, where build-up of CO_2_ and bicarbonate ions during breath-hold diving or digestion facilitates O_2_ delivery to tissues to sustain aerobic metabolism. At the same time, Hb desaturation facilitates CO_2_ transport in the blood as protein-bound CO_2_ and bicarbonate. Corroborating these conclusions, a recent study on caiman blood reports increase in protein-bound CO_2_ and bicarbonate during diving, with bicarbonate binding progressively increasing upon Hb deoxygenation (Bautista et al., 2021). These distinct effects of CO_2_ and bicarbonate on Hb-O_2_ affinity are shared by a phylogenetically diverse set of caiman, alligator, and crocodile species, indicating that these unique regulatory properties evolved prior to the diversification of extant crocodilians >80-100 million years ago in the late Cretaceous, but after the stem lineage of crocodilians diverged from the ancestors of dinosaurs and modern birds.

## Competing Interest Statement

The authors declare no conflict of interest, financial or otherwise.

## Funding

This research was supported by funding from the National Institutes of Health (HL087216) (J.F.S. and A.F.), the National Science Foundation (OIA-1736249) (J.F.S.), and the Independent Research Fund Denmark (T.W. and H.M.).

## Acknowledgments

We thank Elin E. Petersen and Marie Skou Pedersen, for their technical assistance. We also would like to thank Rene Hedegaard and the staff from the Krokodille Zoo (Eskilstrup, Denmark) for assistance in blood sampling. We also thank Vikas Kumar (Mass Spectrometry and Proteomics Core Facility, University of Nebraska Medical Center) for assistance with the MS/MS data analysis.

## Notes

### Competing Interest Statement

The authors have declared no competing interest.

